# Estimating the rate and determinants of exclusive breastfeeding practices among rural mothers in Southern Ghana

**DOI:** 10.1101/543884

**Authors:** Alfred Kwesi Manyeh, Alberta Amu, Rosemond Akpene Ekey, David Etsey Akpakli, John E. Williams, Margaret Gyapong

**Author notes:** Corresponding author P. O. Box. DD1, Dodowa Accra.

## Abstract

**Background:** The health benefits of exclusive breastfeeding practices in both short and long term go beyond the breastfed infant. The benefits of exclusive breastfeeding practices accrue to mothers, families and the society at large. Despite the evidence of these benefits and adoption of various WHO strategies on promotion of exclusive breastfeeding by Ghana, the increase in the rate of exclusive breastfeeding has been very slow in the country. This study aimed to estimate the rate and investigate socioeconomic and demographic determinants of exclusive breastfeeding in two rural districts in Southern Ghana.

**Methods:** Pregnancy, childbirth, breastfeeding, demographic and socioeconomic information of 1,870 women prospectively registered by the Dodowa Health and Demographic Surveillance System and gave birth between January 1, 2011 and December 31, 2013 was extracted. The rate of exclusive breastfeeding among the study participants was estimated and the relationship between the dependent and the independent variables were explored using logistics regression model at 95% confidence level. All data analyses were done in STATA version 14.2.

**Results:** The overall exclusive breastfeeding rate in the study is 70.96 %. Mothers aged 25-29 and 30+ years are 93 and 91 % respectively more likely to practice exclusive breastfeeding compared to those aged <20 years (OR:1.93, 95%CI: 1.25-2.99, OR: 1.91, 95%CI: 1.91-3.08). The odds of artisans practicing exclusive breast feeding is 64% less likely compared to those unemployed (OR: 0.64, 95%CI:0.43-0.96). There is a high chance that 45% of mothers with a household size of more than five members to practice exclusive breastfeeding compared to those with household size of less than six (OR:1.45, 95%CI:1.16-1.81). There is reduced odds of 15% for women in fishing districts compared to those from farming districts (OR:0.15, 95%CI: 0.12 −0.20).

**Conclusion:** There is high rate of exclusive breastfeeding in the study area. Maternal age, type of occupation, household size and district of residence are determinants of exclusive breastfeeding among the study participants.

## Background

The association between child health and development outcomes with appropriate breastfeeding practices such as early initiation at birth, exclusive breastfeeding during the first six months of life, and breastfeeding for at least two years has been established [1, 2]. A baby’s diet during the first few months of life has a significant role on the composition and stability of the gut microbiome that is only acquired after birth [3]. These bacteria make digestion of solids easier, thus, preventing gut problems and illnesses at later stage of life [3]. The World Health Organization (WHO) endorsed exclusive breastfeeding as an optimal way to feed infants [4]. Exclusive breastfeeding is defined by WHO as feeding an infant exclusively with breast milk for the first six months of life. During this period, the infant is allowed to take drops of vitamins, minerals and Oral Rehydration Solution, if prescribed [4].

The health benefits of exclusive breastfeeding practices in both short and long term go beyond the breastfed infant. The benefits of exclusive breastfeeding practices accrue to mothers, families and the society at large [5]. Early suckling enhances the release of prolactin, which helps in the production of milk, and oxytocin, which causes the ejection of milk through the let-down reflex [5]. It also causes contraction of the uterus after childbirth, constricting blood vessels and thereby lessening the likelihood of postpartum hemorrhage [5]. An enhanced breastfeeding practice is known to have reduced child morbidity and mortality, improved quality of life, and enriched human capital [1, 2]. Children who are breastfed correctly for a longer period are known to have a reduced risk of allergies, bowel problems, obesity and diabetes later in adult life, have reduced risk of dental malocclusion, and have higher acumen (children and adolescents) than those who are breastfed for shorter period [2, 6]. Breastfeeding is also associated with positive maternal outcomes such as reduced risk of breast and ovarian cancer, diabetes, and increased birth spacing. It also nurtures mother to child attraction which strengthens the bond between mothers and their babies. In countries where child survival and growth are often endangered by infectious diseases and malnutrition, the benefits of enhanced breastfeeding practices cannot be over emphasized [7-10]. Exclusive breastfeeding is known to subdue the mother’s return to fertility. These effects are influenced by both the duration and regularity of breastfeeding and the age at which the child receives foods and liquids to complement breast milk [5].

Exclusive breastfeeding practices are influenced by multiple factors. These include health, psychosocial, cultural, social, and economic factors [11, 12]. Studies have shown that the decisions regarding exclusive breastfeeding in low-income countries are influenced by education, employment, place of delivery, family pressure, and cultural values [13-16].

Other studies have shown that mixed feeding is associated with increased diarrhea and pneumonia/respiratory diseases in children [17-21]. Infants who were not exclusively breastfed have a 165% higher risk of suffering from diarrhea and 107% higher risk of pneumonia than children who were exclusively breastfed[18, 21]. Worldwide, mixed feeding is attributed to cause the death of 823,000 children under five years of age and 20,000 deaths due to breast cancer in women each year [2].

Despite the evidence on benefits of exclusive breastfeeding, in 2016 only 43% of infants were exclusively breastfed globally [22]. The rate is lower (37%) in low and middle income countries [2]. Despite high child mortality and malnutrition in Sub Saharan Africa (SSA), only 36% of the infants were exclusively breastfed in 2016 [8, 22, 23].

Although Ghana adopted various WHO strategies to promote exclusive breastfeeding, there has been a slow increase in the rates of exclusive breastfeeding in the country. The 2011 Ghana Multiple Indicator Cluster Survey (MICS) in 2011 reported that less than half (46%) of all infants aged 0–6 months in Ghana were exclusively breastfed [24]. This level is lower than that recommended by WHO/UNICEF.

In Ghana, fifty-two percent of children younger than 6 months were exclusively breastfed in 2014 [5]. According to the 2014 Ghana Demographic and Health Survey (GDHS) the percentage of children 0-5 months who were exclusively breastfed has decreased by 17 percent between 2008 and 2014 [5]. The percentage of children who were bottle fed appears to have increased over the past decade. In 2003 and 2014, 11% and 16% of children under 6 months, respectively, were bottled fed [5]. The 2014 GDHS shows that the country still faces the challenges of high infant mortality of 41 deaths per 1,000 live births and 19% of children were stunted ascribable to malnutrition and infections [5].

Using an open cohort longitudinal population-based dataset, we estimated the rate of exclusive breastfeeding examined socioeconomic and demographic factors influencing exclusive breastfeeding practices in two rural districts in Southern Ghana.

## Methods

### Study area and data source

This study was conducted in the Shai-Osudoku and Ningo-Prampram districts of the Greater Accra Region of Ghana. The two districts cover a total population of 115, 754 individuals living in 380 communities in 23,647 households [25]. A comprehensive description of the study districts and the operations of Dodowa Health and Demographic Surveillance System (DHDSS) can be found elsewhere [26-28]. Health service delivery in the study districts is provided by government hospitals, health centres, clinics, Community-based Health and Planning Services (CHPS) compounds/zones, missions and non-governmental health facilities [25, 26]. Secondary longitudinal population-based data was extracted from the database of the DHDSS. The extracted data was exported to STATA version 14.2 for cleaning, coding and analysis.

### Study population

All women resident in the two study districts who were registered in the DHDSS and gave birth between January 1, 2011 and December 31, 2013 were included in the study. Women who were not registered in the DHDSS and those who delivered before 1 January 2011 or after 31 December 2013 were excluded from the study. A total of 1,870 mothers were included.

## Variables

### Dependent variable

The dependent variable is breastfeeding and it was coded as 1 for exclusive breastfeeding and 0 if otherwise. Exclusive breastfeeding in this study is defined as feeding infants with only breast milk, without supplemental liquids or solids except for liquid medicine and vitamin or mineral supplements for the first six months of life [5]. This is based on UNICEF and WHO recommendation [29].

### Independent variables

The independent variables included are maternal age, educational level, marital status, parity, timing of ANC visit, place of delivery, educational level of household head, district of residence and socio-economic status.

The socio-economic status is estimated using weights derived from principal component analysis (PCA) through household social status, ownership of assets, availability of utilities among others [25, 26]. Household socioeconomic status is a proxy measure of a household’s long term standard of living [26]. The proxies from the PCA were divided into five quintiles; poorest, poorer, middle, richer and richest [25, 26].

## Statistical methods

A descriptive analysis of socio-demographic characteristics of the participants was carried out. The rate of exclusive breastfeeding among the study participants was estimated and the relationship between the dependent and the independent variables were explored using logistics regression model. The independent variables that were significant at p < 0.05 in the crude logistics regression model were entered together into an adjusted model. All data analysis were done using Stata version 14.2 and the results were presented in tables with summary statistics at 95% confidence intervals (CI).

## Results

### Socio-Demographic Information

Table 1 presents the socio-demographic information of study participants. The mean age was 27.89 years (SD=7.15).

**Table 1:** Socio-demographic characteristics of the study participants.

| **Characteristics** | **Frequency\*** | **Proportion (%)** |
| --- | --- | --- |
| **Age group** |  |  |
| < 20 | 231 | 12.35 |
| 20-24 | 424 | 22.67 |
| 25-29 | 483 | 25.83 |
| 30+ | 732 | 39.14 |
| **Mean=27.89 (sd=7.15)** |  |  |
| **Ethnicity** |  |  |
| Ga-Dangme | 1,361 | 72.78 |
| Akan | 102 | 5.45 |
| Ewe | 287 | 15.35 |
| Northern | 106 | 5.67 |
| Other tribes | 14 | 0.75 |
| **Religion** |  |  |
| Christianity | 1,706 | 91.23 |
| Islamic | 119 | 6.36 |
| Traditional | 28 | 1.5 |
| Other religions | 17 | 0.91 |
| **Occupation** |  |  |
| Unemployed | 388 | 20.75 |
| Farmer | 307 | 16.42 |
| Artisan | 219 | 11.71 |
| Trader | 594 | 31.76 |
| Civil servant | 33 | 1.76 |
| Student | 289 | 15.45 |
| Others | 40 | 2.14 |
| **Level of education** |  |  |
| No education | 693 | 37.06 |
| Primary | 284 | 15.19 |
| Junior high school level | 656 | 35.08 |
| Senior high school level and above | 237 | 12.67 |
| **Marital status** |  |  |
| Single | 543 | 29.46 |
| Married | 382 | 20.73 |
| Separated/divorced | 36 | 1.9 |
| Cohabiting | 882 | 47.86 |
| **Sex of household head** |  |  |
| Female | 730 | 39.04 |
| Male | 1,140 | 60.96 |
| **Household size** |  |  |
| Less than six | 956 | 51.12 |
| More than five | 914 | 48.88 |
| Mean=6.45 (sd=4.33) |  |  |
| **District of Residence** |  |  |
| Shai-Osudoku | 836 | 44.71 |
| Ningo-Prampram | 1,034 | 55.29 |
| **Parity** |  |  |
| Parity 1 | 520 | 27.81 |
| Parity 2 | 429 | 22.94 |
| Parity 3 | 374 | 20.00 |
| Parity3+ | 547 | 29.25 |
| **Delivery place** |  |  |
| Health facility | 1,332 | 71.23 |
| Outside health facility | 538 | 28.77 |
| **Timing of ANC initiation** |  |  |
| First trimester | 525 | 28.1 |
| Second trimester | 1,196 | 64.03 |
| Third trimester | 147 | 7.89 |
| **Breastfeeding type** |  |  |
| Not exclusive | 543 | 29.04 |
| Exclusive | 1,327 | 70.96 |
| **Exclusive breastfeeding rate** | **70.96%** |  |

*n* = 1870; *SD* = Standard Deviation \* number of respondents across some categories may not add up to 1870 due to missing data

Participants aged <20 years formed the smallest percentage of the study participants (12.35%), while the 30+ age group contributed the highest percentage (39.14%) followed by the 25-29 and the 20-24 years’ groups which accounted for 25.85% and 22.67 % respectively. Majority of the study participants (72.78%) were of the Ga-Dangme ethnic group and 15.35% were Ewes. A large proportion (91.23%) of the participants were of the Christian faith while 6.36% and 1.50% were Moslem and Traditionalist respectively.

A total of 31.76 % of the participants were petty traders, 20.75 % and 16.42 % were unemployed and farmers respectively. Students formed 15.45 % of the study participants. Participants with no formal education and Junior high school level contributed 37.06 % and 35.08 % respectively.

A significant percentage of the study participants (47.86%) were cohabiting with their partners. More than half (60.96%) of the participants’ households were headed by males. The mean household size is 6.45 with a standard deviation of 4.33.

Most of the study participants (55.29%) resided in the Ningo-Prampram District.

Participants with parity 3 or more formed 29.25 % of the study population while those with parity 1 and 2 were 27.81, and 22.94 % respectively.

A large percentage of 71.23 of study participants delivered in a health facility. More than half (64.03%) of the study participants initiated antenatal clinic visit in the second trimester of their last pregnancy.

The overall exclusive breastfeeding rate during the study period was 70.96 %.

### Unadjusted and adjusted odds ratio of determinants of exclusive breastfeeding

Table 2 presents the unadjusted and adjusted Odds Ratio (OR) at 95% Confidence Interval (CI) of socioeconomic and demographic determinants of exclusive breastfeeding in the Dodowa Health and Demographic Surveillance site.

**Table 2:**
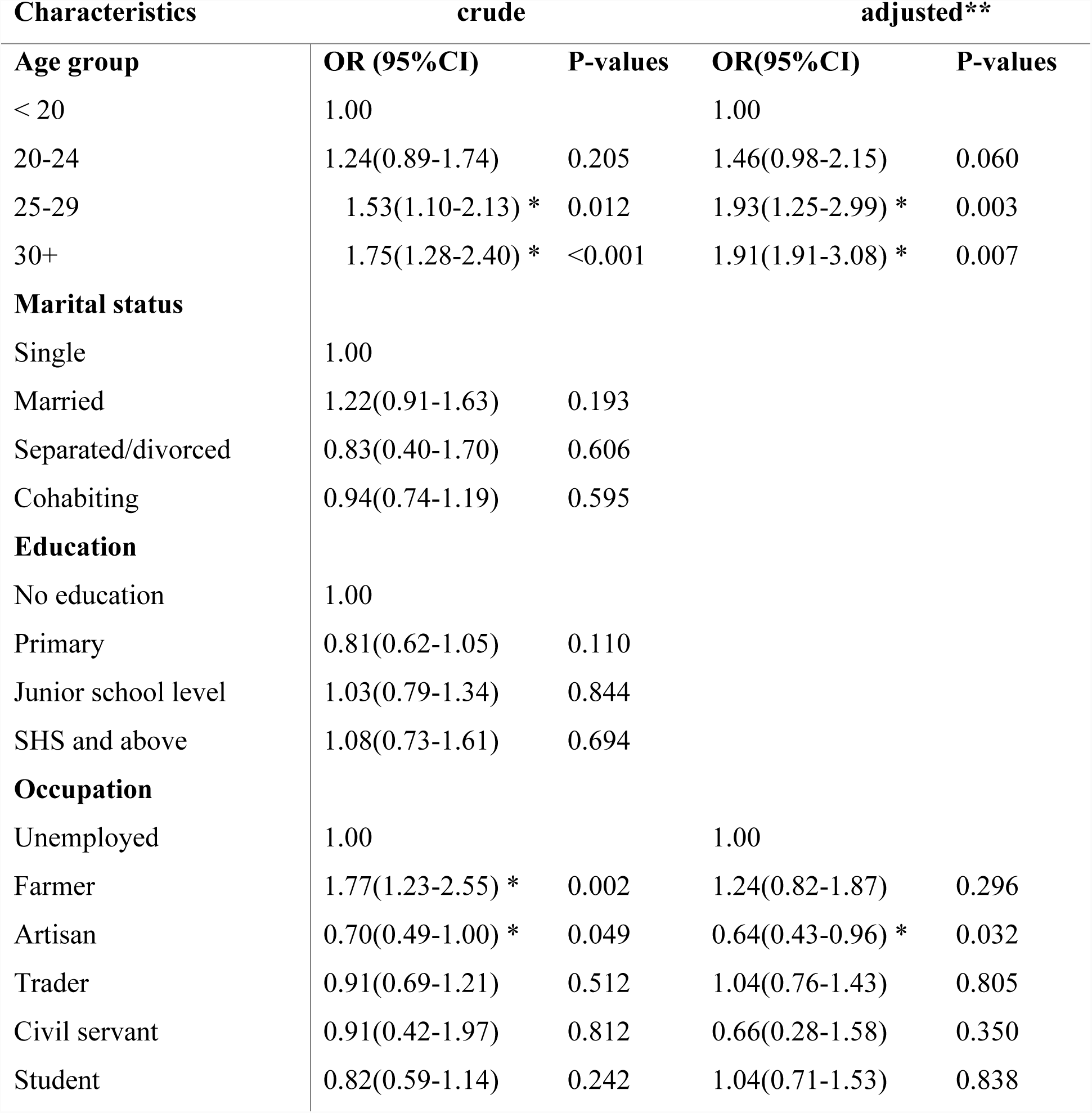
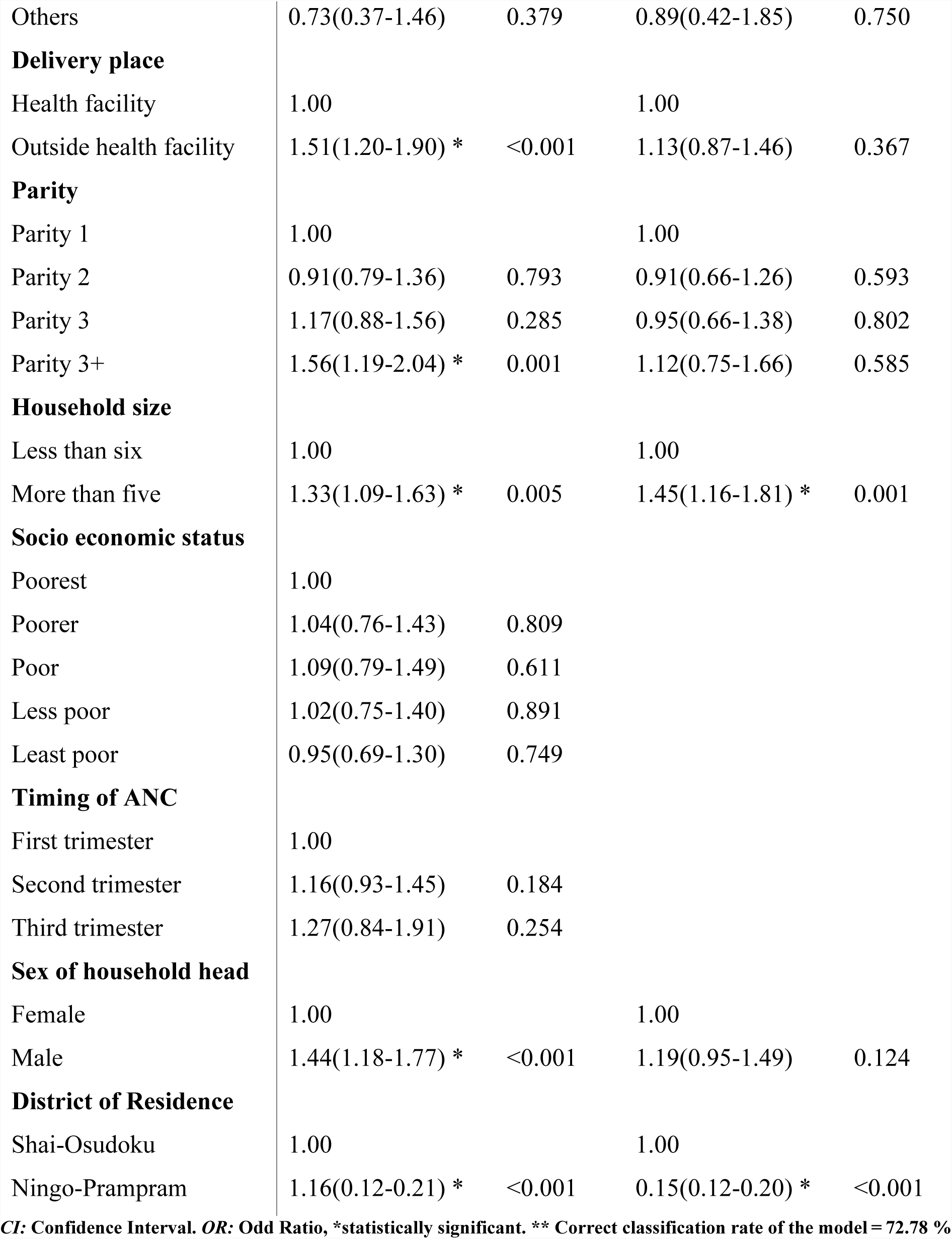
Unadjusted and adjusted odds ratios of factors of exclusive breastfeeding.

| **Characteristics** | **crude** |  | **adjusted\*\*** |  |
| --- | --- | --- | --- | --- |
| **Age group** | **OR (95%CI)** | **P-values** | **OR(95%CI)** | **P-values** |
| < 20 | 1.00 |  | 1.00 |  |
| 20-24 | 1.24(0.89-1.74) | 0.205 | 1.46(0.98-2.15) | 0.060 |
| 25-29 | 1.53(1.10-2.13) * | 0.012 | 1.93(1.25-2.99) * | 0.003 |
| 30+ | 1.75(1.28-2.40) * | <0.001 | 1.91(1.91-3.08) * | 0.007 |
| **Marital status** |  |  |  |  |
| Single | 1.00 |  |  |  |
| Married | 1.22(0.91-1.63) | 0.193 |  |  |
| Separated/divorced | 0.83(0.40-1.70) | 0.606 |  |  |
| Cohabiting | 0.94(0.74-1.19) | 0.595 |  |  |
| **Education** |  |  |  |  |
| No education | 1.00 |  |  |  |
| Primary | 0.81(0.62-1.05) | 0.110 |  |  |
| Junior school level | 1.03(0.79-1.34) | 0.844 |  |  |
| SHS and above | 1.08(0.73-1.61) | 0.694 |  |  |
| **Occupation** |  |  |  |  |
| Unemployed | 1.00 |  | 1.00 |  |
| Farmer | 1.77(1.23-2.55) * | 0.002 | 1.24(0.82-1.87) | 0.296 |
| Artisan | 0.70(0.49-1.00) * | 0.049 | 0.64(0.43-0.96) * | 0.032 |
| Trader | 0.91(0.69-1.21) | 0.512 | 1.04(0.76-1.43) | 0.805 |
| Civil servant | 0.91(0.42-1.97) | 0.812 | 0.66(0.28-1.58) | 0.350 |
| Student | 0.82(0.59-1.14) | 0.242 | 1.04(0.71-1.53) | 0.838 |
| Others | 0.73(0.37-1.46) | 0.379 | 0.89(0.42-1.85) | 0.750 |
| **Delivery place** |  |  |  |  |
| Health facility | 1.00 |  | 1.00 |  |
| Outside health facility | 1.51(1.20-1.90) * | <0.001 | 1.13(0.87-1.46) | 0.367 |
| **Parity** |  |  |  |  |
| Parity 1 | 1.00 |  | 1.00 |  |
| Parity 2 | 0.91(0.79-1.36) | 0.793 | 0.91(0.66-1.26) | 0.593 |
| Parity 3 | 1.17(0.88-1.56) | 0.285 | 0.95(0.66-1.38) | 0.802 |
| Parity 3+ | 1.56(1.19-2.04) * | 0.001 | 1.12(0.75-1.66) | 0.585 |
| **Household size** |  |  |  |  |
| Less than six | 1.00 |  | 1.00 |  |
| More than five | 1.33(1.09-1.63) * | 0.005 | 1.45(1.16-1.81) * | 0.001 |
| **Socio economic status** |  |  |  |  |
| Poorest | 1.00 |  |  |  |
| Poorer | 1.04(0.76-1.43) | 0.809 |  |  |
| Poor | 1.09(0.79-1.49) | 0.611 |  |  |
| Less poor | 1.02(0.75-1.40) | 0.891 |  |  |
| Least poor | 0.95(0.69-1.30) | 0.749 |  |  |
| **Timing of ANC** |  |  |  |  |
| First trimester | 1.00 |  |  |  |
| Second trimester | 1.16(0.93-1.45) | 0.184 |  |  |
| Third trimester | 1.27(0.84-1.91) | 0.254 |  |  |
| **Sex of household head** |  |  |  |  |
| Female | 1.00 |  | 1.00 |  |
| Male | 1.44(1.18-1.77) * | <0.001 | 1.19(0.95-1.49) | 0.124 |
| **District of Residence** |  |  |  |  |
| Shai-Osudoku | 1.00 |  | 1.00 |  |
| Ningo-Prampram | 1.16(0.12-0.21) * | <0.001 | 0.15(0.12-0.20) * | <0.001 |

CI: Confidence Interval. OR: Odd Ratio, \*statistically significant. \*\* Correct classification rate of the model = 72.78 %

In the unadjusted model, maternal age, occupation, place of delivery, parity, household size and sex of household head had statistically significant associations with exclusive breastfeeding.

The odds of mothers aged 20-24 practicing exclusive breastfeeding is 24% more likely compared to those aged <20 years (OR: 1.24, 95%CI: 0.89-1.74). Women aged 25-29 and 30+ years are 53% and 75% more likely to practice exclusive breastfeeding respectively compared to those aged <20 years (OR: 1.53, 95%CI: 1.10-2.13, OR: 1.75, 95%CI: 1.28-2.40). This is statistically significant. A similar pattern is observed in the adjusted model such that in the presence of other explanatory variables, the odds of mothers practicing exclusive breastfeeding increased with increasing maternal age. Mothers aged 20-24 years were 46% more like to practice exclusive breastfeeding compared to those aged <20 years (OR: 1.46, 95%CI: 0.98-2.15). The odds of mothers aged 25- 29 and 30+ years are 93 and 91 % respectively more likely to practice exclusive breastfeeding compared to those aged <20 years (OR:1.93, 95%CI: 1.25-2.99, OR: 1.91, 95%CI: 1.91-3.08). This is statistically significant.

Mothers who were married were 22% more likely to practice exclusive breastfeeding compared to those who were single (OR: 1.22, 95%CI: 0.91-1.63). Participants who were separated/divorced and cohabiting have a reduced odds of 83% and 94% respectively of practicing exclusive breastfeeding compared to those who were single (OR: 0.83, 95%CI: 0.40-1.70, OR:0.94, 95%CI: 0.74-1.19).

The results further revealed that, the odds of a mother practicing exclusive breast feeding is 81% less likely associated with a mother with primary level of education compared to those with no education (OR:0.81, 95%CI: 0.62-1.05). Again, the odds of women with Junior and senior level of school practicing exclusive breastfeeding is 3% and 8% (respectively) more likely compared to those with no education (OR: 1.03, 95%CI: 0.79-1.34, OR:1.08, 95%CI:0.73-1.61).

There was a statistically significant association between occupation and women practicing exclusive breastfeeding. The odds of a farmer practicing exclusive breastfeeding is 77% more likely compared to those unemployed (OR: 1.77, 95%CI: 1.23-2.55). Women artisans were 70% less likely to practice exclusive breastfeeding compared to those unemployed (OR: 0.70, 95%CI: 0.49-1.00). Mothers who were traders, civil servants and students were 91%, 91% and 82% respectively less likely to practice exclusive breastfeeding compared to those unemployed (OR:0.91, 95%CI:0.69-1.21, OR:0.91, 95%CI:0.42-1.97, OR: 0.82, 95%CI: 0.59-1.14). In the adjusted model, farmers, traders and students are 24%, 4% and 4% more likely to feed their babies exclusively with breast milk compared to women unemployed (OR: 1.24, 95%CI: 0.82-1.87, OR: 1.04, 95%CI: 0.76-1.43, OR:1.04, 95%CI: 0.71-1.53). The odds of female artisans and civil servants to practice exclusive breast feeding is less likely compared to those unemployed (OR: 0.64, 95%CI:0.43-0.96, OR:0.66, 95%CI:0.28-1.58).

In the unadjusted model, the place of delivery was significantly associated with mothers practicing exclusive breastfeeding such that, the odds of women who delivered outside health facility to practice exclusive breastfeeding is 51% more likely compared to those who delivered in a health facility (OR:1.51, 95%CI: 1.20-190). This association was not statistically significant (OR:1.13, 95%CI: 0.87-1.46) in the adjusted model.

In the crude model, the odds of women with parities 2 to practice exclusive breastfeeding compared to those with parity 1 was 0.91 (OR: 0.91, 95%CI: 0.79-1.36). There was an increased odds of 17% and 56% of mothers with parities 3 and more (OR: 1.17, 95%CI: 0.88-1.56, OR:1.56, 95%CI: 1.19-2.04). In the presence of other variables (Age, Occupation, Place of delivery, Household size, Sex of household head and District of residence), while there was reduced odds of 91% and 95% of women with parities 2 and 3 respectively to practice exclusive breastfeeding (OR:0.91, 95%CI:0.66-1.26, OR:0.95, 95%CI:0.66-1.38), there was increased odds of 12% for mothers with parity more than 3 (OR:1.12, 95%CI: 0.75-1.66) in the adjusted model.

There is an increased odds of 33% and 45% in the crude and adjusted model respective of mothers with household size more than five members to practice exclusive breastfeeding compared to those with household size of less than six (OR: 1.33, 95%CI:1.09-1.63, OR:1.45, 95%CI:1.16-1.81). This was also statistically significant.

In the crude analysis, participants who belong to poorer, poor and less poor socioeconomic categories were 4%, 9% and 2% respectively more likely to practice exclusive breastfeeding compared to those in poorest category.

There was increased odds of 16% and 27% for participants who initiated antenatal visit in the second and third trimesters respectively to practice exclusive breastfeeding compared to those who initiated theirs in the first trimester (OR: 1.16, 95%CI:0.93-1.45, OR:1.27, 95%CI:0.84-1.91). The crude analysis showed a significant association of women whose households were headed by males being 44% more likely to practice exclusive breastfeeding compared to those in female headed households (OR: 1.44, 95%CI: 1.18-1.77).

The district of residence of participant is statistically significantly associated with practicing exclusive breastfeeding; there was an increased odds of 16% for women from Ningo-Prampram district compared to those from Shai-Osudoku district in the unadjusted analysis. The odds ratio rather reduced with the adjusted model to 0.15 ((OR:1.16, 95%CI: 0.12-0.21, unadjusted) (OR:0.15, 95%CI: 0.12.0.20, adjusted)). This is statistically significant.

## Discussion

The aim of this study was to estimate the rate and examine socioeconomic and demographic determinants of exclusive breastfeeding in two rural districts (Ningo-Prampram and Shai-Osudoku Districts) of Southern Ghana.

The exclusive breastfeeding rate among the study participants in this study is 70.96%. This rate is higher than 52% reported by 2014 GDHS [5] but lower than 84.3% report by another Ghanaian study [30].

Maternal age, type of occupation, household size and district of residence appear to be strong determinants of exclusive breastfeeding practices after adjusting for other variables.

Older women were more likely to practice exclusive breastfeeding compared to younger mothers. This result is consistent with studies in other settings which showed that, younger mothers are at an increased risk of early cessation of exclusive breastfeeding[31-35]. Our finding is also similar to a study conducted in Tanzania where participants who were younger were less likely to practice exclusive breastfeeding [36].

Although maternal level of education and socioeconomic status were found elsewhere to be significantly associated with breastfeeding practices in other studies [37], the current study has not found a statistically significant association between maternal education, socioeconomic status and exclusive breastfeeding.

This supports the finding of an earlier study [38] which suggested that educational attainment of mothers was not associated with exclusive breastfeeding. Nonetheless, the finding of this current study contradicts the results of studies in Nigeria [39, 40], Tanzania [36], India [41, 42], Brazil [38] and in Ghana [24] which reported that the level of educational attainment of mothers was positively associated with exclusive breastfeeding practice. Due to the education provided to pregnant women at highly patronized antenatal care (87%) [5], the issue of exclusive breastfeeding has become a universal knowledge hence it is not preserved for only educated women in the study area.

The high rate of exclusive breastfeeding among the study participants can also be attributed to education from Reproductive and Child Health (RCH) Centers of the Ghana Health Service in the study area where pregnant women receive antenatal care services with education on breastfeeding practices as shown in other study [30].

The findings show that, mothers who are self-employed artisans are more likely to practice exclusive breastfeeding. This finding is in line with earlier studies that found self-employed mothers to be more likely to practice exclusive breastfeeding [43]. This study found that mothers whose households were headed by males were more likely to practice exclusive breastfeeding but this relationship is not statistically significant after adjusting for other explanatory variables. This result is similar to the findings of other studies in Ghana [43] and Malawi [44] where the decision of exclusive breastfeeding is influenced by spouses and family members.

The relationship between household size and exclusive breastfeeding as found in this study has also been established in another study [45]. Mothers with family size of 4 and less were more likely to practices exclusive breastfeeding as compared those with family size above 4 [45].

It was also very intriguing to find that the district of residence is significantly associated with exclusive breastfeeding practice among the study participants. Women from Ningo-Prampram District are 15% less likely to practice exclusive breast feeding compared to those from Shai-Osudoku District. This can be ascribed to the introduction of pregnancy schools in health facilities in Shai-Osudoku District where expectant mothers are educated on how to handle themselves and their babies and the importance of exclusive breastfeeding among others. The effective RCH in Shai-Osudoku District where pregnant women receive antenatal care services with education on breastfeeding practices could be another contributing factor to the high likelihood of exclusive breast feeding practice in the Shai-Osudoku District.

## Strength and limitations of the study

Despite the advantage of large sample and use of population based data, this study has a number of limitations. First, the outcome was measured based on self-report; recall bias may have been underestimated or overestimated the association between the outcome of interest and the explanation variables. Social desirability bias could also be a limitation to the study as some women might have withheld what they thought to be negative aspects of their breastfeeding practices. The study did not explore other factors such as knowledge, initiation, duration and cultural determinants of exclusive breastfeeding which might have some influence on the outcome of interest. This is primarily due to the limited information in the secondary data used. The study was also limited to only two districts in the Greater Accra Region of Ghana hence, limits the generalizability of the findings.

## Conclusions

Majority of the study participants practiced exclusive breastfeeding. Maternal age, type of occupation, household size and district of residence are strong determinants of exclusive breastfeeding practices.

Maintaining access to information on appropriate breastfeeding practices and promotion of exclusive breastfeeding especially among young mothers in the study area is highly recommended. To further understand other factors influencing the practice of exclusive breastfeeding and to design a suitable evidence-based intervention targeting young mothers, we recommend further qualitative study in this area.

## Abbreviations

ANC: : Antenatal Care
CI: : Confidence Intervals
DHDSS: : Dodowa Health and Demographic Surveillance System
DHRC: : Dodowa Health Research Centre
GDHS: : Ghana Demographic Health Survey
JHS: : Junior High school
OR: : Odd Ratio
PCA: : Principal component analysis
RCH: : Reproductive and Child Health
WHO: : World Health Organization

## Acknowledgments

We sincerely thank the study participants, and staff of the DHDSS for their continuous support to the HDSS. This study is dedicated to Prof. Margaret Gyapong who worked tirelessly to establish the Dodowa Health and Demographic Surveillance System.

## Availability of data and materials

All relevant data supporting the conclusions of this article are included within the article.

## Authors’ contributions

AKM conceptualized, designed, conducted data extraction and the statistical analysis for the study. He also led the drafting of the paper. AA, RAE and DEA contributed to the initial design of the study, literature review and drafting of the paper. JW and MG refined the study design and critically reviewed the paper. All authors read and approved the manuscript.

## Consent to participate and Ethics approval

At the beginning of each data collection round, Dodowa Health Research Centre sought verbal consent from household heads and all individual participants as shown in earlier studies [25, 26]. The Ethical Committee of Ghana Health Service and the Institutional Review Board of Dodowa Health Research Centre approved the operations, data collection procedure and quality assurance of the DHDSS [25, 26]. The management of Dodowa Health Research Centre sanctioned the use of data for this study on a condition that the participants’ identity remains anonymous.

## Consent for publication

Not applicable.

## Reference

1. Rollins NC, Bhandari N, Hajeebhoy N, Horton S, Lutter CK, Martines JC, et al: Why invest, and what it will take to improve breastfeeding practices? Lancet 2016, 387(10017):491–504.

2. Victora CG, Bahl R, Barros AJD, França GVA, Horton S, Krasevec J: Breastfeeding in the 21st century: epidemiology, mechanisms, and lifelong effect. Lancet 2016, 387(10017):475–490.

3. Sullivan AO, Farver M, Smilowitz JT: The Influence of Early Infant-Feeding Pratices on the Intestinal Microbiome and Body Composition in Infants. Nutrition and Metabolic Insights 2015, 8(1):87.

4. UNICEF, WHO: Indicators for assessing infant and young child feeding practices. In: Part 1 Definitions. Geneva: WHO; 2008.

5. Ghana Statistical Service (GSS), Ghana Health Service (GHS), ICF International: Ghana Demographic and Health Survey 2014. In. Rockville, Maryland, USA: GSS, GHS, ICF International; 2015.

6. Victora CG, Horta BL, de Mola CL, Quevedo L, Pinheiro RT, Gigante DP, et al: Association between breastfeeding and intelligence, educational attainment, and income at 30 years of age: a prospective birth cohort study from Brazil. Lancet Global Health 2015, 3(4):e199–e205.

7. Jones G, Steketee RW, Black RE, Bhutta ZA, Morris SS: How many child deaths can we prevent this year?. Lancet 2003, 362(9377):65–71.

8. Black RE, Alderman H, Bhutta ZA, Gillespie S, Haddad L, Horton S, Lartey A, Mannar V, Ruel M, Victora CG et al: Maternal and child nutrition: Building momentum for impact. Lancet 2013, 382:372–375.

9. Black RE, Victora CG, Walker SP, Bhutta ZA, Christian P, de Onis M, et al: Maternal and child undernutrition and overweight in low-income and middle-income countries. Lancet 2013, 382(9890):427–451.

10. Black RE, Allen LH, Bhutta ZA, Caulfield LE, de Onis M, Ezzati M, et al: Maternal and child undernutrition: global and regional exposures and health consequences. Lancet 2008, 371(9608):243–260.

11. Cripe ET: Supporting breastfeeding(?):nursing mothers’ resistance to and accommodation of medical and social discourses. In Emerging Perspective in Health Communication: Meaning, Culture and Power. New York: Routledge Taylor and Francis Group; 2008.

12. Schmied V, Barclay L: Connection and pleasure, disruption and distress: Women’s experience of breastfeeding. J Hum Lact 1999, 15(4):325–334.

13. Gartner LM, Morton J, Lawrence RA, Naylor AJ, O’Hare D, Schanler RJ, Eidelman AI: American Academy of Pediatrics Section on Breastfeeding: Breastfeeding and the use of human milk. Pediatrics 2005, 115:496–506.

14. Ogunlesi TA: Maternal socio-demographic factors influencing the initiation and exclusivity of breastfeeding in a Nigerian semi-urban setting. Matern Child Health J 2010, 14(3):459–465.

15. Henry BA, Nicolau AI, Americo CF, Ximenes LO, Bernheim RG, Oria MOB: Socio-cultural factors influencing breastfeeding practices among low-income women in Fortaleza-Ceara-Brazil: a Leininger’s sunrise model perspective. Enfermeria Global 2010.

16. Otoo GE, Lartey AA, Pérez-Escamilla R,. -: Perceived incentives and barriers to exclusive breastfeeding among Periurban Ghanaian women. J Hum Lact 2009, 25(1):34–41.

17. Horta B, Victora C: Short-term Effects of breastfeeding: a systematic review on the benefits of breastfeeding on diarrhoea and pneumonia mortality. World heal Organ 2013:1–54.

18. Lamberti LM, Zakarija-grkovi I, Walker CLF, Theodoratou E, Nair H, Campbell H, et al: Breastfeeding for reducing the risk of pneumonia morbidity and mortality in children under two: a systematic literature review and metaanalysis. BMC Public Health 2013, 13:s18.

19. Hanieh S, Ha TT, Simpson JA, Thuy TT, Khuong NC, Thoang DD, et al: Exclusive breast feeding in early infancy reduces the risk of inpatient admission for diarrhea and suspected pneumonia in rural Vietnam: a prospective cohort study. BMC Public Health 2015, 15:1166.

20. Nkemjika SO, Demissie K: Breast feeding initiation time and its impact on diarrheal disease and pneumonia in West Africa. J Public Heal Epidemiol 2015, 7:352–359.

21. Lamberti LM, Walker CLF, Noiman A, Victora C, Black RE: Breastfeeding and the risk for diarrhea morbidity and mortality. BMC Public Health 2011, 11:S15.

22. Mgongo M, Hussein TH, Stray-Pedersen B, Vangen S, Msuya SE, Wandel M: “We give water or porridge, but we don’t really know what the child wants:” a qualitative study on women’s perceptions and practises regarding exclusive breastfeeding in Kilimanjaro region, Tanzania. BMC Pregnancy and Childbirth 2018, 18:323.

23. Yalcin SS, Berde AS, Yalcin S, ;. Determinants of exclusive breastfeeding in sub Saharan Africa: a multilevel approach. Paediatr Perinat Epidemiol 2016, 30(5):439–449.

24. Ghana Statistical Service: Ghana Multiple Indicator Cluster Survey with an Enhanced Malaria Module and Biomarker. In. Accra Ghana; 2011.

25. Manyeh AK, Amu A, Akpakli AE, Williams J, Gyapong M: Socioeconomic and demographic factors associated with caesarean section delivery in Southern Ghana: evidence from INDEPTH Network member site. BMC Pregnancy and Childbirth 2018, 18:404.

26. Manyeh AK, Kukula V, Odonkor G, Ekey RA, Adjei A, Narh-Bana S, Akpakli DE, Gyapong M: Socioeconomic and demographic determinants of birth weight in southern rural Ghana: evidence from Dodowa health and demographic surveillance system. BMC Pregnancy and Childbirth 2016, 16:160.

27. Awini E, Sarpong D, Adjei A, Manyeh AK, Amu A, Akweongo P, Adongo P, Kukula V, Odonkor G, Narh S et al: Estimating cause of adult (15+years) death using InterVA-4 in a rural district of southern Ghana. Global Health Action 2014, 7(1):25543.

28. Gyapong M, Sarpong D, Awini E, Manyeh KA, Tei D, Odonkor G, Agyepong IA, Mattah P, Wontuo P, Attaa-Pomaa M et al: Health and demographic surveillance system profile: the Dodowa HDSS. Int J Epidemiol 2013(42):1686–1696.

29. Begin F, Arts M, White J, Clark D, Sint TT, Taqi I, et al: From the First Hour of Life - Making the Case for Improved Infant and Young Child Feeding Everywhere. In. New York: UNICEF; 2016.

30. Boakye-Yiadom A, Yidana A, Sam NB, Kolog B, Abotsi A: Factors Associated with Exclusive Breastfeeding Practices among Women in the West Mamprusi District in Northern Ghana: A Cross-Sectional Study. 2016, 6(3):91–98.

31. Avery M., Duckett L., Dodgson J., Savik K., Henly S.J: Factors associated with very early weaning among primiparas intending to breastfeed. Matern Child Health J 1998, 2:167–179.

32. Hauck Y.L., Fenwick J., Dhaliwal S.S., Butt J.: A Western Australian survey of breastfeeding initiation, prevalence and early cessation patterns. Matern Child Health J 2011, 15:260–268.

33. Liu P., Qiao L., Xu F., Zhang M., Wang Y., Binns C.W.: Factors Associated with Breastfeeding Duration. J Hum Lact 2013, 29:253–259.

34. Kaneko A., Kaneita Y., Yokoyama E., Miyake T., Harano S., Suzuki K., Ibuka E., Tsutsui T., Yamamoto Y., Ohida T.: Factors associated with exclusive breast-feeding in Japan: For activities to support child-rearing with breast-feeding. J Epidemiol 2006, 16:57–63.

35. Ludvigsson J.F., Ludvigsson J.: Socio-economic determinants, maternal smoking and coffee consumption, and exclusive breastfeeding in 10205 children. Acta Paediatr 2005, 94:1310–1319.

36. Victor R, Baines SK, Agho KE: Determinants of breastfeeding indicators among children less than 24 months of age in Tanzania: a secondary analysis of the 2010 Tanzania Demographic and Health Survey BMJ oPEN 2013, 3:e001529.

37. Aidam BA, Pérez-Escamilla R, Lartey A, Aidam J: Factors associated with exclusive breastfeeding in Accra, Ghana. European Journal of Clinical Nutrition 2005, 59:789–796

38. Masarenhas ML, Singh KT, Sukhwinder S: Prevalence of Exclusive Breast Feeding and its determinants in the first 3 months of life in South of Brazil. Journal of Paediatrics 2006, 82:289–294.

39. Ogbonna C, Okolo AA, Ezeogu A: Factors influencing exclusive breastfeeding in Jos, Plateau State, Nigeria. West Afr J Med 2000, 19:107–110.

40. Aghaji MN: Exclusive breastfeeding practice and associated factors in Enugu Nigeria. West Afr J Med 2002, 21:66–69.

41. Patel A, Badhoniya N, Khadse S, et al: Infant and young child feeding indicators and determinants of poor feeding practices in India: secondary data analysis of National Family Health Survey 2005–06. Food Nutr Bull 2010, 31:314–333.

42. Rajesh K, Panna CP, Abhay BK: Breast Feeding Initiation Practice and Factors Affecting Breast Feeding in South-Guajat Region of Indian. Journal of Family Practice 2009.

43. Mensah KA, Acheampong E, Anokye FO, Okyere P, Appiah-Brempong E, Adjei RO: Factors influencing the practice of exclusive breastfeeding among nursing mothers in a peri-urban district of Ghana. BMC Research Notes 2017, 10:466.

44. Kent CJ, Mitoulas RL, Cregan DM, Ramsay TD, Doherty AD, Hartmann EP: Volume and frequency of breastfeedings and fat content of breast milk throughout the day Pediatrics 2006, 117(3):e387–395.

45. Reddy S, Abuka T: Determinants of Exclusive Breastfeeding Practice among Mothers of Children Under Two Years Old In Dilla Zuria District, Gedeo Zone, Snnpr, Ethiopia, 2014. J Preg Child Health (2016, 3:224.

